# Simultaneous spike-time locking to multiple frequencies

**DOI:** 10.1101/149906

**Authors:** Fabian H. Sinz, Carolin Sachgau, Jörg Henninger, Jan Benda, Jan Grewe

**Author notes:** Corresponding author: Fabian Sinz.

## Abstract

Phase locking of neural firing is ubiquitously observed in the brain and occurs when neurons fire at a particular phase of a periodic signal. Here we study in detail how spikes of single neurons can simultaneously lock to multiple distinct frequencies at the example of p-type electroreceptor afferents in the electrosensory system of the Gymnotiform weakly electric fish *Apteronotus leptorhynchus*. We identify key elements for multiple frequency locking, study its determining factors and limits, and provide concise mathematical models reproducing our main findings. Our findings provide another example how rate and temporal codes can coexist and complement each other in single neurons, and demonstrate that sensory coding in p-type electroreceptor afferents provides a much richer representation of the sensory environment than commonly assumed. Since the underlying mechanisms are not specific to the electrosensory system, our results could provide the basis for studying multiple-frequency locking in other systems.

## Introduction

Phase locking in neuronal activity is observed when neurons fire action potentials at or around a particular phase of a periodic signal. It is a widespread phenomenon in neural systems and has been studied intensively in the auditory (Köppl, 1997; Surlykke et al., 1988; Joris and Smith, 2008; Kayser et al., 2012, 2009), mechano-sensory (Taniguchi and Hisashu, 1987), electrosensory (Bastian, 1981; Chacron et al., 2000; Ratnam and Nelson, 2000; van Hemmen et al., 2011), visual (Montemurro et al., 2008; Martin and Schröder, 2016; Belitski et al., 2008), vibrissial (Ewert et al., 2008), and the lateral line system (Wubbels, 1992). The phase of spikes can carry important information about the timing of external signals or provide additional information in time-scale multiplexed neural codes (Panzeri et al., 2010; Bullock, 1997; Lisman, 2005; Belitski et al., 2008; Kayser et al., 2009). However, most studies focus on locking to a single frequency and the connection of the locking to single cell properties remains elusive.

Action potentials recorded in p-type electroreceptor afferents of gymnotiform wave-type weakly electric fish are well known to phase-lock to their self-generated, quasi-sinusoidal electric field, called the electric organ discharge (EOD, Fig. 1 A–C) (Hagiwara and Morita, 1963; Bastian, 1981). Similar to the noticeable modulation in volume when listening to two almost identical tones, the superposition of two EODs during social interactions causes a periodic amplitude modulation (AM) of the electric field, called beat, which oscillates at the absolute difference between the single EOD frequencies (Fig. 1 D). Amplitude modulations are encoded in p-type electrorecptor afferents, so called P-units (Scheich et al., 1973; Bastian, 1981; Benda et al., 2005; Walz et al., 2014), that are distributed over the body of the fish (Carr et al., 1982). These neurons usually fire at most once per EOD period and vary their spiking probability per EOD cycle with the amplitude of the EOD (Scheich et al., 1973; Knudsen, 1974; Hopkins, 1976; Benda et al., 2005). Because their peri-stimulus time histogram (PSTH) at EOD resolution closely resembles the shape of the amplitude envelope of their field, P-units are thought of as AM encoders and their response properties have mainly been characterized in terms of the AM of the EOD (Fig. 1 D) (Wessel et al., 1996; Chacron et al., 2003; Grewe et al., 2017). In the absence of AMs P-units also encode phase modulations of a periodic carrier (Carlson and Kawasaki, 2006).

**Figure 1.**
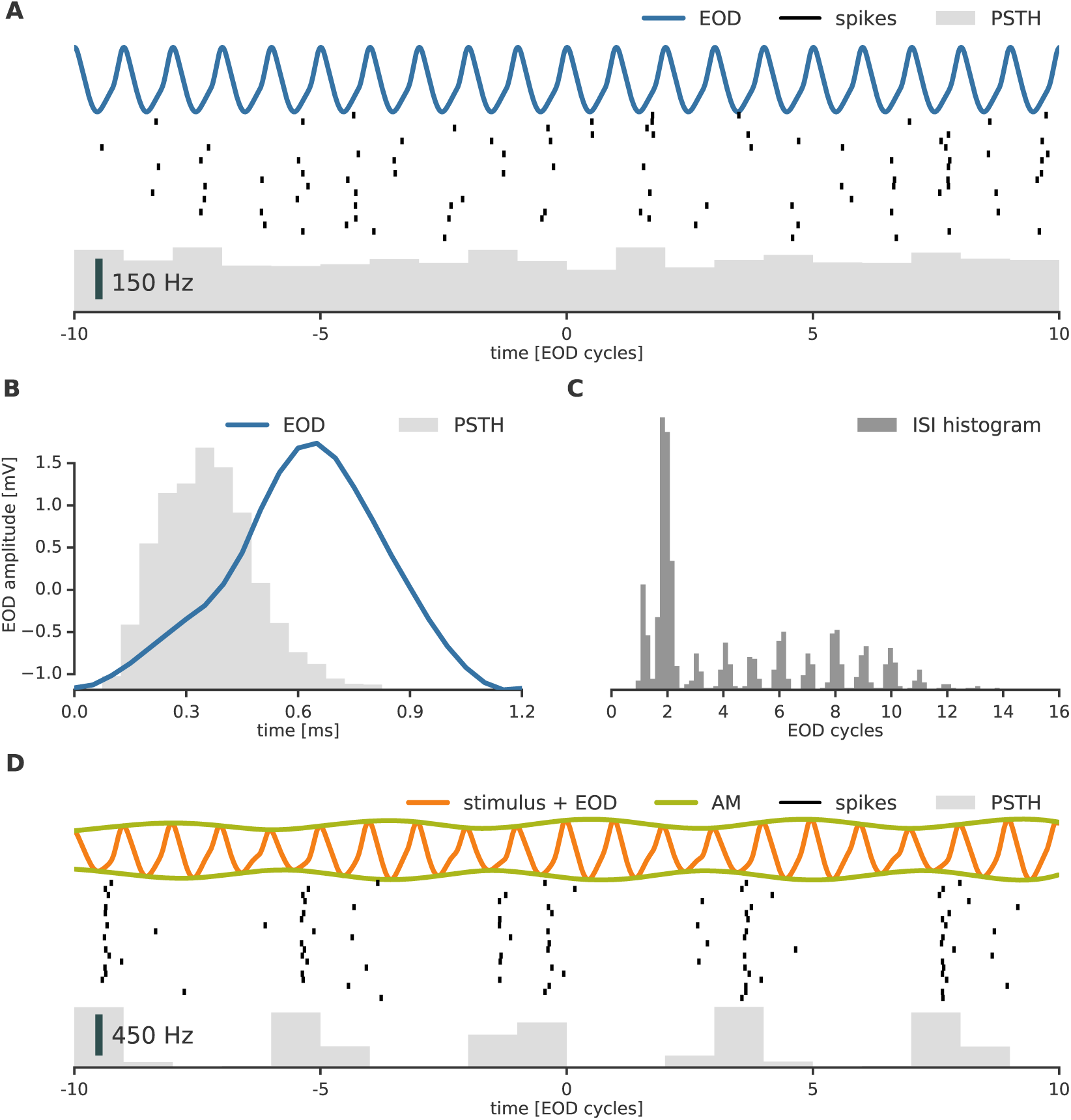
Locking to the EOD in an p-type electroreceptor afferent. (cell 2014-12-03-ao) **A** The EOD (top) of Gymnotiform weakly electric fish is a periodic but not sinusoidal signal. Spikes (middle) of p-type electroreceptor afferents fire at high rates phase locked to the EOD (see also B) but not necessarily every cycle (see also C). In the absence of an external stimulus, the PSTH binned at EOD cycle resolution is uniform (bottom). **B** Cyclic histogram (bars) of baseline spiking for one EOD period: Within each EOD cycle (solid line) electroreceptor afferents preferentially fire to a particular phase of the EOD. Note that the relative alignment of the PSTH and the EOD is arbitrary since it contains an unknown delay because electric field travel almost instantaneous while neuronal signals take some time to reach the recording site. **C** Multi-modal inter-spike-interval (ISI) histogram of the baseline activity of a single electroreceptor afferent. Cycle skipping causes peaks to occur on multiples of the EOD period. **D** An external periodic signal generates an amplitude modulation (AM, green) of the EOD (orange). This causes the P-units to modulate their firing rate with the AM. At EOD resolution, the firing rate modulation follows the AM of the external field (bottom). As in B, there is an arbitrary relative delay between AM and the spikes. PSTHs in A and D were computed from large sets to temporal segments of the cellular response, raster plots depict only a small subset (20 segments) of these.

In their natural neotropical habitats, weakly electric fish are exposed to a wide range of frequencies from conspecific and heterospecific EODs (Stamper et al., 2012). Since EOD frequencies of individuals from the same species are relatively close (Crampton and Albert, 2006), intra-specific interactions usually cause low-frequency AMs of at maximum a few hundred Hertz (Walz et al., 2014), while cross-species interactions yield beat AMs of much higher frequencies. In addition to the relative frequency encoded by the AM, the absolute frequency of an external EOD might be of behavioral relevance for the fish. However, whether information about the absolute frequency is encoded in electroreceptor afferents has not been studied yet.

Here, we demonstrate that p-type electroreceptor afferents in *Apteronotus leptorhynchus* provide a rich representation of multiple sine-wave stimuli in their rate as well as their spike timing. Not only do spikes lock to the frequency of the resulting periodic AM, but also directly to each of the frequencies of the presented sinusoidal signals over a range of several hundred Hertz. We explore the origin, the limits, and ambiguities of this locking to multiple frequencies, and provide a simple leaky integrate-and-fire (LIF) model that can qualitatively reproduce the empirical observations. Finally, we show that multiple locking also occurs in pyramidal cells in the electrosensory lateral line lobe (ELL, Maler, 1979), the next stage of electrosensory processing. The mechanisms and mathematical models we explore here are not specific to the electrosensory system, and we hope that our results also generate new insights in other systems such as, for instance, the closely related auditory system.

## Results

In order to mimic the fields of other weakly electric fish, we presented sine wave stimuli of various frequencies and amplitudes to *n* = 4 specimen of *A. leptorhynchus* while recording intracellular activity from *n* = 7 p-type electroreceptor afferents in the anterior lateral line nerve or the ELL (see Methods and Materials). P-units were identified by several features: a baseline firing rate of 64–470 Hz (Ratnam and Nelson, 2000; Gussin et al., 2007; Grewe et al., 2017), a multi-modal interspike-interval (ISI) histogram (Fig. 1 C) caused by the locking to a specific phase of the EOD (Fig. 1 B), and a sigmoidal firing rate vs. input curve (*f*-*I*-curve) estimated from the response to amplitude steps in the EOD field.

### P-units lock to a wide range of stimulus frequencies

P-units represent stimulus information at two time scales: timing information originating from skipping EOD periods at a fine scale of less than one EOD cycle (Fig. 1 A–C), and amplitude information at a coarse scale of one EOD cycle (Fig. 1 D). In the presence of an external signal, the firing rate follows the timecourse of the resulting AM. At the same time, locking to the EOD creates an additional temporal structure within each AM period (Fig. 2 A). If P-units were simple amplitude encoders, spiking at random times within each EOD period would suffice. The rather precise firing within the EOD (Fig. 1 B; Fig. 2 A) in unison with the rate modulation induced by the AM makes the timing of P-unit spikes carry information as well.

We measured spike timing information using vector-strength spectra (van Hemmen et al., 2011). The vector strength 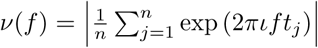 quantifies locking of *n* spike times *t_j_* to a particular frequency *f*. *ι* denotes the imaginary unit. Intuitively, it computes the mean of all spike times placed according to their phase 2*πft_j_* on a unit circle whose circumference represents one period 1/*f* of a signal with frequency *f*. If the spikes are spread symmetrically around the entire circle then *ν* = 0, if they perfectly lock to the stimulus they cluster all on one phase and *ν* = 1, otherwise 0 < *ν* < 1. We computed the vector-strength spectrum averaged over trials for each combination of stimulus frequency and amplitude and refer to these spectra as second order. In contrast, first order spectra were computed by accumulating spikes over trials before the vector strength was calculated (see Methods and Materials). Second order spectra preserve locking to both the applied stimulus and the ongoing EOD of the fish, while first order spectra potentially loose locking to the EOD but are less prone to sampling noise due to low spike numbers.

In all recorded P-units, we found locking not only to the EOD, but also to the beat, as well as harmonic combinations of those (locking defined as *ν*(*f*) significantly different from zero with *α* = 0.001; see Methods and Materials). To our surprise, we additionally found direct locking to the absolute stimulus frequency even when it was several hundred Hertz above the EOD frequency which would typically be considered to be outside the range of P-unit tuning (Knudsen, 1974; Hopkins, 1976) (Fig. 3). For instance, the cell shown in Fig. 2B exhibits a clear locking to a stimulus with a frequency well above 1000 Hz (peak at *f_s_* = 1368 Hz).

**Figure 2.**
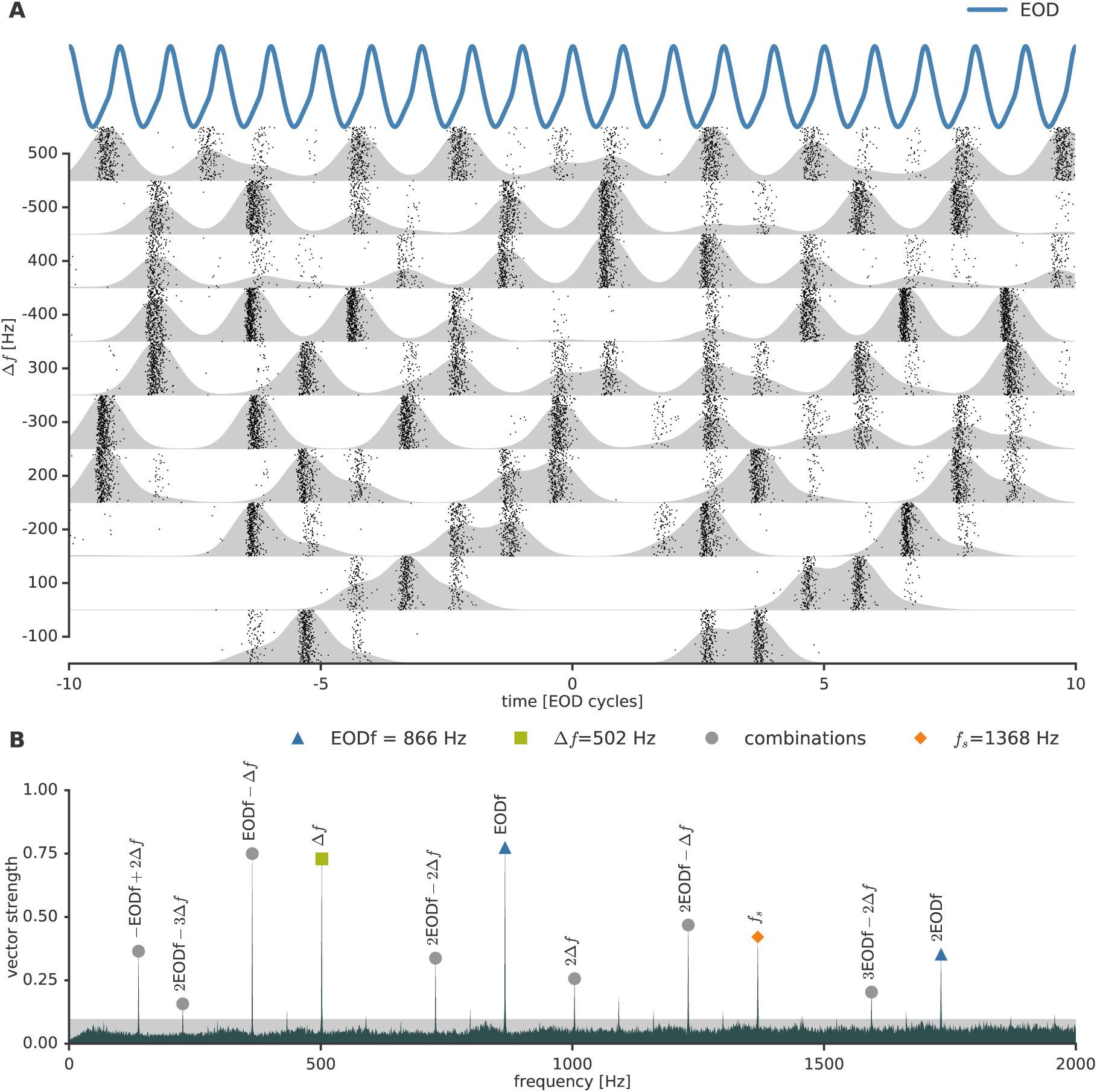
Locking to the stimulus frequency in a P-unit. (cell 2014-12-03-ao) **A** Spike-raster plot of a P-unit stimulated with sine wave stimuli of different frequencies. Δ*f* specifies the difference between the stimulus frequency and the EOD frequency and is the frequency of the resulting beat AM. The firing probability (shaded regions; PSTH low-pass filtered with Gaussian with *σ* ≈ 0.5ms) is modulated with the AM frequency Δ*f*. Spike rasters are aligned to zero phase difference between peaks in EOD (top) and stimulus (*t* = 0; stimulus not shown). Because of locking to a particular phase within each EOD period, there is local temporal structure within each period of the AM while, at the same time, firing rates are modulated by the AM on a coarser time scale. **B** Second order vector-strength spectrum of spike trains elicited by a 20% contrast stimulus Δ*f* = 502 Hz above the EOD frequency. The height of the shaded area depicts critical value for spurious locking at *α* = 0.001 for each frequency (see Methods). The cell clearly exhibits locking at the EOD frequency EOD *f* = 866 Hz (triangle), at the stimulus frequency *f_s_* = EOD*f* + Δ*f* = 1368 Hz (diamond), as well as EOD*f* − Δ*f* and Δ*f* = 502 Hz (square). Other peaks in the spectrum result from per trial higher order interactions of spike times (circles).

**Figure 3.**
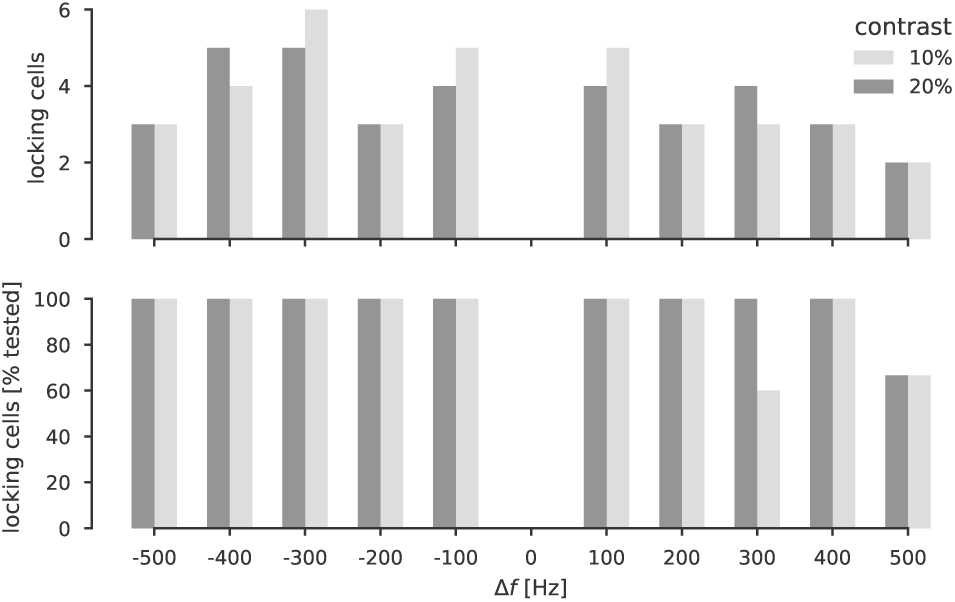
Locking to stimulus frequencies in P-units. Over a wide range of stimulus frequencies almost all recorded P-units showed locking to the absolute stimulus frequency. Locking is defined as a vector strength significantly different from zero (*α* = 0.001; see Methods and Materials). Δ*f* denotes difference between the stimulus frequency and the EOD frequency. Top shows the absolute numbers of cells found locking, bottom shows the number of locking cells in percent of tested cells for that combination of contrast and Δ*f*. The contrast indicates the stimulus amplitude relative to the fish’s EOD amplitude.

### How can a spike train lock to more than one frequency at the same time?

Simultaneous locking to two or more frequencies occurs when the relative distribution of spike times is asymmetric within one period of the locking frequency (Fig. 4 A–B). In P-units this can be achieved through the finely grained temporal structure resulting from the locking to the EOD: In the absence of an external signal, the spike density exhibits a peak at each EOD period (Fig. 4 C, left). When wrapping the spike density around a circle whose circumference (period) does not match the period of the EOD frequency or one of its harmonics, the resulting distribution will be symmetric with respect to the origin (Fig. 4 C, middle). Note, that a period mismatch does not necessarily lead to a uniform distribution because the two periods can have a least common multiple. However, it will still be symmetric, so that the mean of the distribution on the plane will still be at zero (marked with a black dot in Fig. 4 C, middle). Since locking is the vector norm of that mean, there is no locking in that situation.

**Figure 4.**
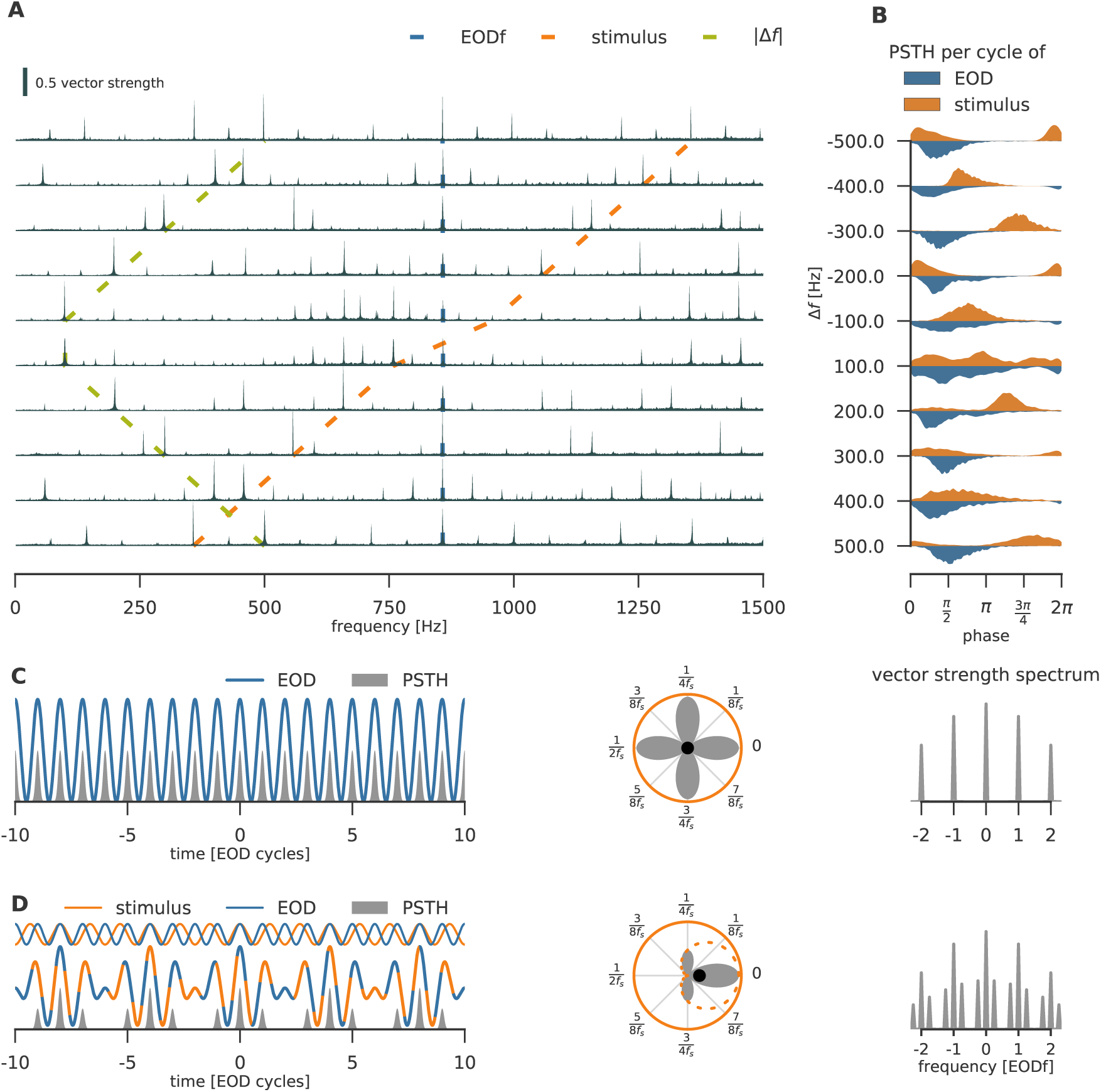
Mechanisms of locking. (cell: 2014-12-03-aj) **A** Second order vector strength spectrum *ν*(*f*) for different stimulus frequencies *f* and corresponding AM frequencies Δ *f* as indicated in B (rows). The external field amplitude was set to 20% of the EOD amplitude (20% contrast). The electroreceptor afferent clearly exhibits locking to the stimulus frequency (orange), the EOD frequency (blue), and the absolute difference |Δ*f* | between the two (green) over a wide range of frequencies. Other peaks result from higher order interactions of spike times. **B** Cyclic spike densities relative to single period of the stimulus (upper) and the EOD (lower) for the same stimulus frequencies as in A. The single mode in each density indicates locking to that particular frequency (colors as in A). **C** Left: Illustrative schematic of the spike density (gray) evoked by the EOD (blue line) without an external stimulus. Middle: When wrapping the spike density around a circle whose circumference is the period 1/*f_s_* of a putative stimulus (here *f_s_* = 3/4 · EOD *f*) the resulting density is symmetric with respect to the origin, the center of mass is at the origin and there is no locking to this frequency. Right: Corresponding vector-strength (Fourier) spectrum of the baseline spike density. **D** Left: Illustrative schematic of the spike density (gray) in the presence of a superposition (blue-orange dashed) of the EOD and the stimulus (blue and orange, respectively; shown not to scale), creating an AM with a period of 4 EOD cycles. The AM reaches its peak when the peak of the stimulus and the EOD coincide. Middle: The stimulus (orange, dashed) acts as a weighting function on the spike density, shifting the center of mass (vector strength) of the density away from zero (round marker), thereby creating locking. Since locking to the EOD is still preserved, this neurons now locks to two frequencies. Right: Vector strength (Fourier) spectrum of the spike density.

When the cell is driven by an external periodic stimulus, however, the spike density will be modulated with the absolute value of the frequency difference Δ*f* between the stimulus and the EOD (Fig. 4 D, left), because the AM and thus the firing rate is enhanced when signal and EOD are in phase and suppressed when they are out of phase. Thus, the stimulus signal effectively becomes a weighting function for the PSTH peaks (dashed line in Fig. 4 D, middle). When wrapped around a circle whose circumference is equal to the stimulus period, all peaks of the spike density on one side of the circle, where the AM amplitude is high, will be enhanced. Note that the AM amplitude can only be high where the stimulus itself is large. All peaks on the other side of the circle will be suppressed, because there the stimulus and thus the AM amplitude is low. This makes the initially symmetric PSTH asymmetric, thereby shifting the center of gravity away from zero, inducing locking to the stimulus frequency.

### Ambiguities in locking

Whenever we found locking to the stimulus frequency EOD*f* + Δ*f*, we also found locking to EOD*f* − Δ*f*. This ambiguity can be explained with a simple phenomenological model of the spike density. The vector strength spectrum is proportional to the Fourier transform of the spike train (Ashida et al., 2010, see eqn. (1) in Methods and Materials). When spikes are aligned to the stimulus and aggregated over trials before computing the spectrum (first order spectrum), the vector strength spectrum converges to the amplitude spectrum of the PSTH. Therefore, we can use the Fourier transform of the PSTH to understand the locking effects in more detail.

The phenomenological model explains locking to the external stimulus — and weaker locking to its harmonics — by locking of the electroreceptor afferent to the EOD and Gaussian jitter on the spike times. It assumes that each electroreceptor afferent fires at a particular phase of the EOD period and that the spike times are jittered by Gaussian random noise. This means that the PSTH is given by regularly spaced Gaussians whose width is determined by the noise variance (Fig. 4 C, left). Using properties of the Fourier transform, this implies that the vector strength spectrum has peaks at multiples of the EOD frequency (EOD*f*) weighted by a Gaussian centered at zero whose variance is inversely proportional to the noise variance (Fig. 4 C, right; see Phenomenological Model in Methods and Materials).

Incorporating the firing rate modulation into the model demonstrates that locking must occur at EOD*f* ± Δ*f* : A simple way to capture the modulation of the firing rate in the presence of a stimulus is to weight (multiply) the periodic spike density with a sine of frequency Δ*f* before jittering the spike times (Fig. 4 D, left). Using properties of the Fourier transform (multiplication in temporal space becomes convolution in frequency space, see Phenomenological Model in Methods and Materials), this causes each peak in the vector strength spectrum to be flanked by two additional peaks at Δ*f* to the left and right (Fig. 4 D, right), as observed in vector strength spectra of our data.

The second order vector strength spectra of spike trains recorded *in vivo* exhibit more peaks than predicted by this simple model (Fig. 4 A). These are caused by nonlinear interactions between the harmonics of the EOD and the stimulus through the spiking mechanism of the neuron (see LIF model below). The real EOD has harmonics, because it is periodic but not exactly sinusoidal. In contrast to taking the Fourier transform of the PSTH (first order spectrum), computing single trial vector strength spectra and averaging those (second order spectrum), preserves single trial spike time dependencies which show up at harmonics of the frequencies involved and linear combinations thereof (see Methods and Materials).

### A leaky integrate-and-fire model can qualitatively account for the main locking effects

Next we investigated whether the main peaks found *in vivo* are created by the non-linear spike mechanism in a more realistic neuron model. To this end, we drove a leaky integrate-and-fire model (LIF) with the rectified and low-pass filtered output of a harmonic oscillator. The oscillator itself was driven by the electric field composed of the EOD and the stimulus (Walz, 2013, see also Fig. 5 and Methods and Materials). In contrast to the phenomenological model above, this model generates spikes as a direct function of the electric field. Apart from setting the resonance frequency of the oscillator to the EOD frequency, the model parameters were not adapted to a particular neuron but were manually adjusted to lead to a generally reasonable model behavior. The amplitude spectrum of the electric field shows three distinct peaks (Fig. 5A). The most prominent peak is located at the EOD frequency and, because the EOD is periodic but not purely sinusoidal, also at the second harmonic. The stimulus, corresponding to the foreign fish’s EOD, leads to a peak at the stimulus frequency. The harmonic oscillator captures the tuning of the receptor organ to the EOD (Hopkins, 1976; Viancour, 1979) and acts as a band-pass filter on the stimulus, attenuating the stimulus peak in the spectrum (Fig. 5 A). Rectification by the receptor synapse (Chacron et al., 2000) and subsequent low-pass filtering by the afferent’s dendrite extracts the amplitude modulation of the EOD and hence introduces an additional peak at the AM frequency as well as an interaction of the AM and the second harmonic of the EOD (Fig. 5 B). Finally, the nonlinear spiking mechanism of the LIF gives rise to the second symmetric peak at EOD*f* − Δ*f* below the EOD frequency and other peaks caused by higher order interactions in the vector strength spectrum (Fig. 5 C). The spectrum of this simple model therefore reproduces the main features of the spectrum of a real electroreceptor afferent faithfully (compare Fig. 5 C and 5 D), including many, but not all, additional peaks in the locking.

**Figure 5.**
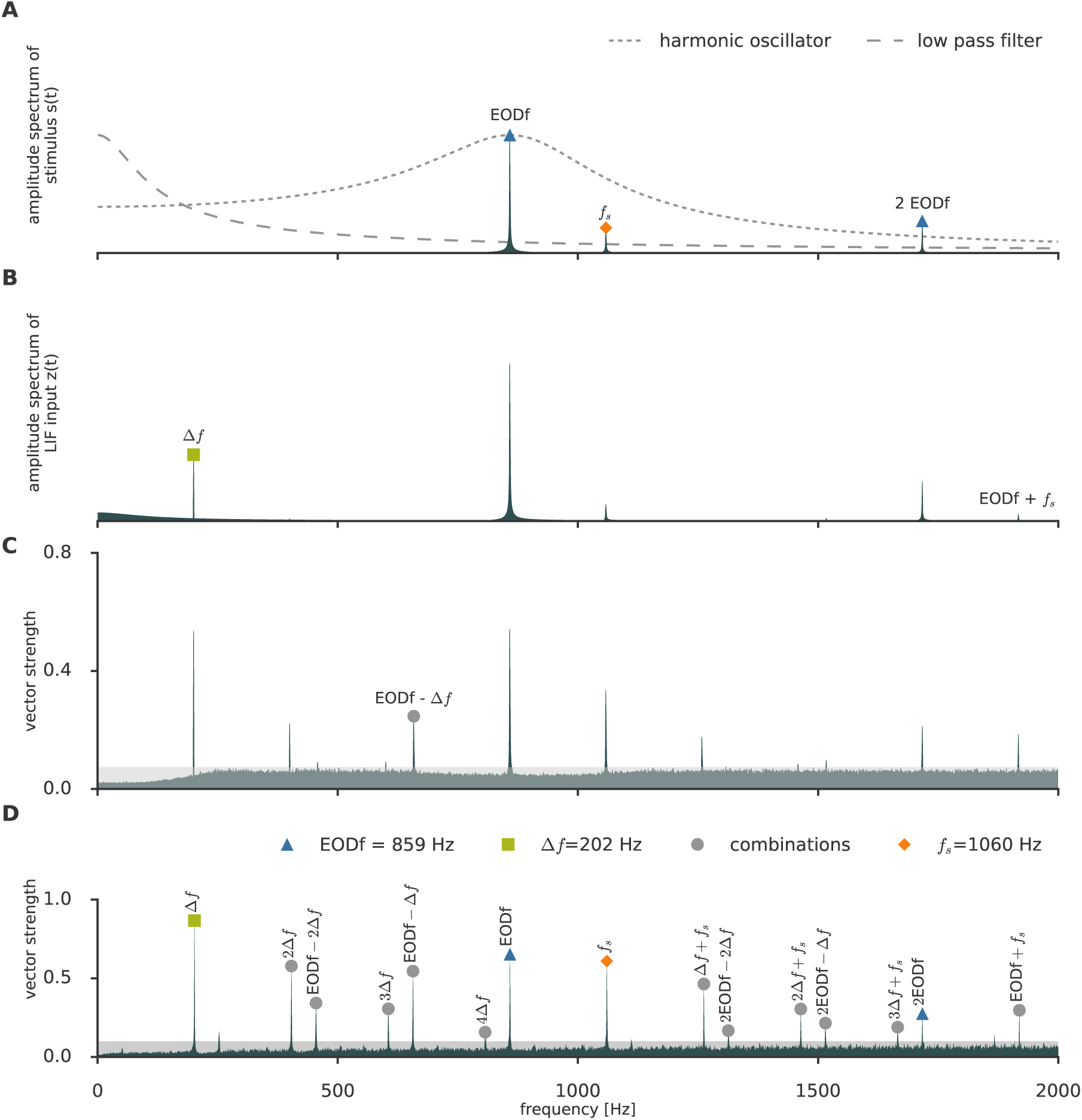
Multiple locking peaks in a leaky integrate-and-fire model. **A** Amplitude spectrum (gray) of the electric field in the presence of an external signal. Because the EOD is not perfectly sinusoidal, it exhibits harmonics (peaks at EOD*f* and 2 · EOD*f*). The initial harmonic oscillator of the electroreceptor acts as a band-pass filter on the spectrum. After rectification at the receptor synapses, a low-pass filter models the passive properties of the afferent dendrite. **B** Amplitude spectrum of the input to the LIF model, which is the rectified and low-pass filtered output of the harmonic oscillator which itself is driven by the signal whose spectrum is shown in A. Rectification and low-pass filtering approximate a Hilbert transform and introduce a peak at the AM frequency (Δ*f*) **C** The signal in B is the input to a LIF model. Second order vector strength spectrum of the spike trains generated by the LIF model. The spiking mechanism introduces the second symmetric peak to the left of the EOD frequency and other harmonics. **D** Example second order spectrum from a real P-unit with the same EOD frequency (cell 2014-12-03-ai). The real spectrum contains several peaks at interactions of the different frequency components of the signal that the model does not account for.

### Factors and limits of locking

From the simple phenomenological model above, we can identify three components that should affect locking: the stimulus frequency, the jitter of spike times within an EOD period, and the amplitude of the external signal (Fig. 6). We quantified the influence of those parameters on the locking in 119 trials, referring to a particular combination of stimulus contrast and frequency, recorded in 7 electroreceptor afferents, where spikes significantly locked on the stimulus frequency (*p* ≤ 0.001 tested per trial; see Methods and Materials for test). As expected, the vector strength is proportional to stimulus amplitude (contrast relative of EOD amplitude, *ρ* = 0.36, *p* < 5 · 10^−5^, Fig. 6 A) — the stronger the stimulus the better the locking — and inversely proportional to the stimulus frequency (*ρ* = −0.64, *p* < 3 · 10^−7^, Fig. 6 B) and the circular standard deviation (*ρ* = −0.40, *p* < 0.003, Fig. 6 C) at a fixed stimulus contrast of 20%. The circular standard deviation of the spike times relative to one EOD period during stimulation, which is an estimator of the standard deviation of a circularly wrapped normal distribution, quantifies spike-time jitter. However, for a given circular standard deviation, locking at higher stimulus frequencies should be more degraded by spike time jitter than locking at lower frequencies (coloring in Fig. 6 C). To quantify the joint influence of the different factors, we fitted a generalized linear model (*γ*-likelihood, inverse link function) to predict the vector strength from all three of them. We also added a cross term between frequency and circular standard deviation. The circular standard deviation and frequency interaction term, the frequency term, contrast term, and the circular standard deviation all show a significant influence on the locking (*p* < 0.01).

**Figure 6.**
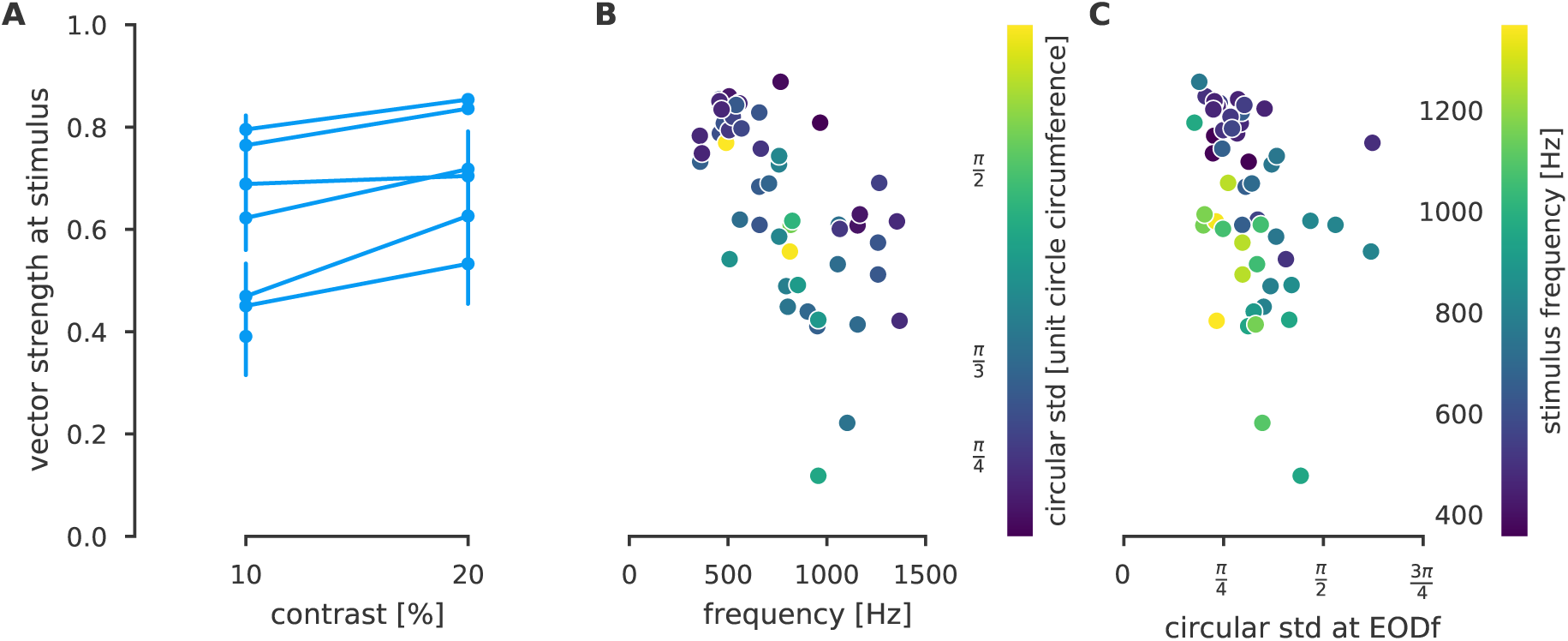
Factors influencing locking to the stimulus frequency. (*n* = 119 trials recorded in *n* = 7 P-units) **A** Vector strength at stimulus frequency as a function of stimulus contrast pooled across all stimulus frequencies (Pearson correlation *ρ* = 0.36, *p* < 5 · 10^−5^). Lines connect contrast measured in the same cell. Points are averages from a single cell. Errorbars depict the 95% confidence interval over trials. **B** Vector strength as a function of stimulus frequency at contrast 20% (Pearson correlation *ρ* = −0.64, *p* < 3 · 10^−7^). Color encodes the circular standard deviation of spike times, i.e. their jitter. **C** Vector strength as a function of circular standard deviation at contrast 20% (Pearson correlation *ρ* = −0.40, *p* < 0.003). Color encodes the stimulus frequency.

The amount of temporal jitter on the spike locking within one EOD period should set an upper limit on the frequencies an electroreceptor afferent can lock to. Following our model considerations from above, the Gaussian temporal jitter translates into a Gaussian weighting profile in the Fourier domain where the temporal standard deviation and the standard deviation in frequency space are related by 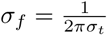. The resulting *σ_f_* cover a range from approximately 350 Hz to 1600 Hz with a mean of about 1060 Hz. This means that the typical EOD frequency range of *A. leptorhynchus* from about 600 Hz to 1000 Hz is well within the 1-sigma range of the low-pass filter on the locking induced by the jitter on the spike times.

### Decoding: Locking to the beat vs. locking to the stimulus

Which signal is better suited to detect the presence of an external periodic field: modulations of the firing rate caused by the AM of the beat induced by the external field or direct locking to the stimulus? In principle, an external sinusoidal signal can be read out from P-unit activity from either source. One way to read out firing rate modulation at particular frequencies from single trials is to compute locking of the spikes to the AM frequency. We therefore quantified locking to the stimulus and the AM frequency for all trials in which we were able to detect significant locking to either signal. In most cases, spike trains lock to both the stimulus and the beat (Fig. 7A, inset). Cases in which the spike train locks solely to the stimulus are rare. We also frequently saw significant locking to the AM frequency even in cases where the stimulus was more than 200 Hz away from the EOD, which is usually considered outside the tuning range of P-units (*n* = 613 out of 622 trials, Fig. 7B). In the vicinity of the EOD, locking to the beat was usually stronger than locking to the stimulus (Fig. 7C). On average, locking to the stimulus was rarely stronger than locking to the beat. beat induced by the external field or direct locking to the stimulus? In principle, an external sinusoidal signal can be read out from P-unit activity from either source. One way to read out firing rate modulation at particular frequencies from single trials is to compute locking of the spikes to the AM frequency. We therefore quantified locking to the stimulus and the AM frequency for all trials in which we were able to detect significant locking to either signal. In most cases, spike trains lock to both the stimulus and the beat (Fig. 7 A, inset). Cases in which the spike train locks solely to the stimulus are rare. We also frequently saw significant locking to the AM frequency even in cases where the stimulus was more than 200 Hz away from the EOD, which is usually considered outside the tuning range of P-units (*n* = 613 out of 622 trials, Fig. 7 B). In the vicinity of the EOD, locking to the beat was usually stronger than locking to the stimulus (Fig. 7 C). On average, locking to the stimulus was rarely stronger than locking to the beat.

**Figure 7.**
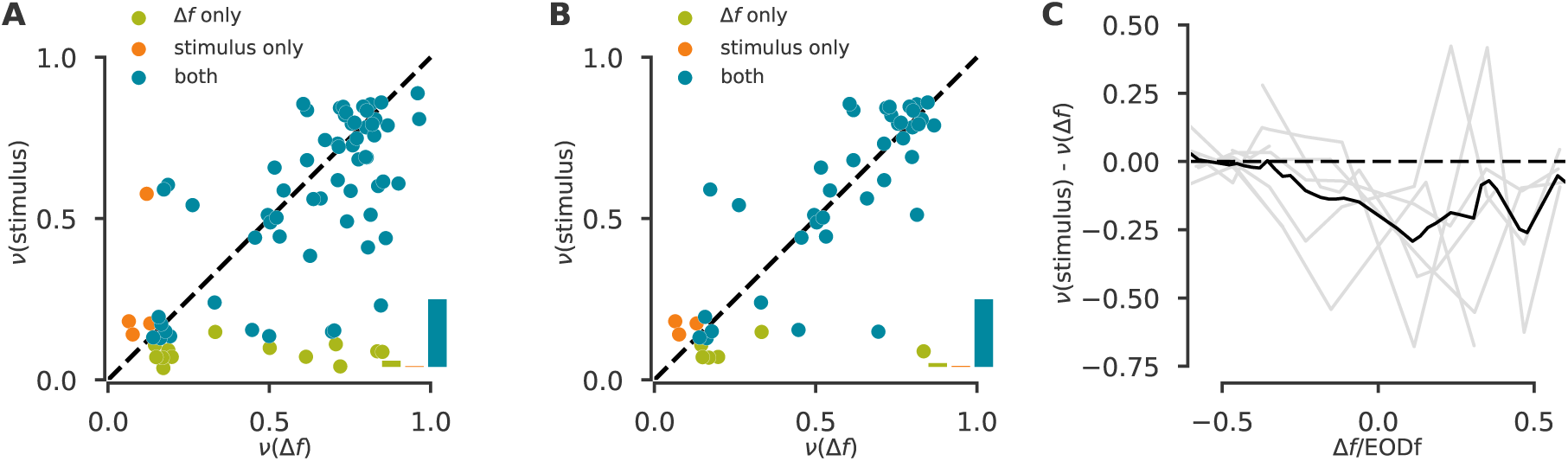
Locking to beat and stimulus. Vector strengths (*ν*) were computed for the beat (Δ*f*) and the stimulus for all trials with 20% contrast in which significant locking to the beat (*n* = 77 trials), the stimulus *n* = 10 trials, or both *n* = 856 was found. **A** Scatter plot of the vector strengths averaged over several trials with the same stimulus parameters. Green points correspond to trials in which only significant locking to the beat was found, orange trials with only locking to the stimulus, and blue where the spike train locked to both. Inset histogram shows the respective proportions. **B** Like in A, but only for trials for which Δ *f >* 200 Hz. There is no noticeable difference in the proportion of trials shown in the insets in A and B. **C** Average difference in vector strength over trials as a function of Δ*f* normalized to the corresponding EOD frequency to put Δ*f* on a common scale. Different lines correspond to different cells. Black line depicts the average over cells. Linear interpolation was used to compute the mean. Points outside the measured range for a given curve were not considered in averaging. P-units seem to lock stronger to the beat than the stimulus (below the diagonal in A) when |Δ*f* | is small. On average, locking to the stimulus is rarely stronger than locking to the beat.

### Downstream processing of spike time locking

P-units project onto pyramidal cell in the electrosensory lateral line lobe (ELL) in the hindbrain which is the only route for all electrosensory information to enter higher brain areas (Maler, 1979; Carr et al., 1982; Maler, 2009). If the electrosensory system uses the locking information provided by P-units, then this information should be present in pyramidal cells as well.

We recorded spikes from *n* = 27 pyramidal cells in the ELL of *n* = 7 specimen of *A. leptorhynchus*. 16 cells showed locking to the external sinusoidal signal (*ν*(*f*) significantly different from zero at *α* = 0.001; see Methods and Materials for test). The characteristics of an example cell are shown in Fig. 8 A–C. Pyramidal cells have interspike-interval histograms that are smoother than those found in P-units (Fig. 8 A) and they exhibit less pronounced locking to the EOD (Fig. 8 B). Since pyramidal cells have a lower firing rate than P-units (25 Hz in this example neuron), second order (per trial) spectra contain substantially more finite sampling noise. We therefore used first order spectra, where we first aligned spike times to the stimulus peaks, accumulated all spikes over trials, and then computed the vector strength spectrum (see Methods and Materials). While this strongly reduces the noise, it can also average out other peaks, apart from the ones from the stimulus. The spectra of the responses to a range of stimulus frequencies is shown in figure 8 C.

**Figure 8.**
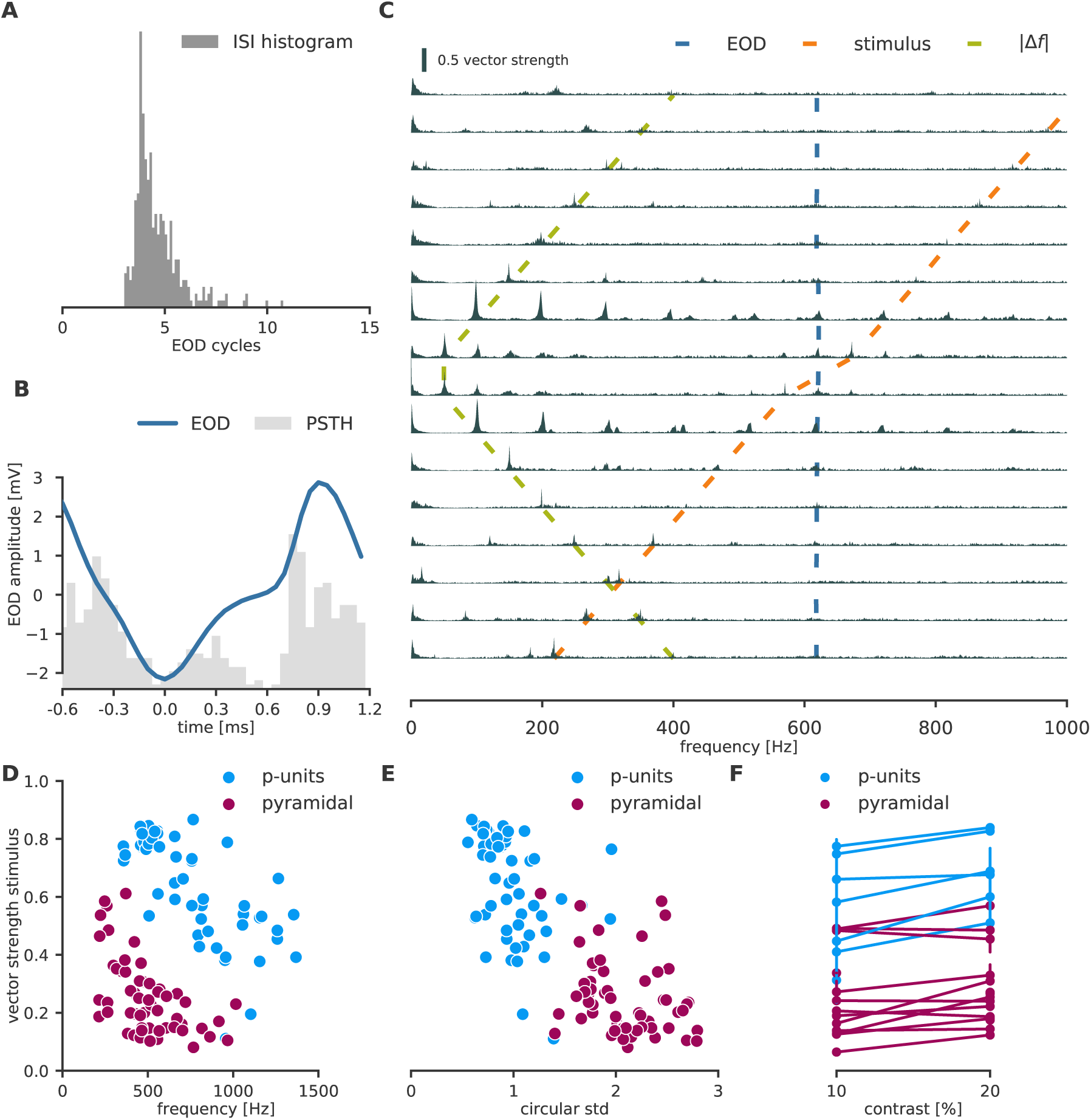
Locking in pyramidal cells. (cell 2014-11-26-ad) **A** The inter-spike interval distribution of pyramidal cells does not show the multimodal structure as the one of p-type electroreceptor afferents. **B** Baseline histogram of spike times for one EOD cycle (blue). The pyramidal cell still locks to the EOD frequency, but less pronounced (vector strength *ν* = 0.23, *p* < 10^−8^, compare to 2 B). **C** First order locking spectra: First, trials were aligned to the stimulus phase and then spikes were aggregated over trials before computing the vector strength spectrum. Therefore, other frequencies than the stimulus might show less locking than on a single trial basis, in particular the EOD frequency. The pyramidal cell exhibits clear locking to the stimulus (orange line) in low frequencies but less in higher frequencies. **D–F** Factors of locking from first order spectra (similar to Figure 6). Locking with respect to frequency and circular standard deviation was computed at contrast 20%. **D** Pyramidal cells show less locking and lock in a smaller frequency range than P-units. **E** Consistent with a lower vector strength, their circular standard deviation of the spike times is larger. In general, locking in pyramidal neurons is weaker than in P-units. **F** Locking in pyramidal cells increases with stimulus amplitude in a similar way as observed in P-units. Error bars represent the 95% confidence interval.

In general, significant locking was weaker in pyramidal neurons than in P-units and was rarely found at high frequencies (Fig. 8 D). However, locking was inversely correlated with stimulus frequency (Pearson correlation *ρ* = −0.5, *p* < 10^−20^) as well as the circular standard deviation (*ρ* = −0.23, *p* < 10^−20^) at 20% contrast. Compared to P-units, the temporal jitter of the spikes per EOD period was also markedly higher (Fig. 8 E). Like in P-units, locking in pyramidal cells was contrast dependent (*ρ* = 0.18, *p* < 10^−20^, Fig. 8 F).

## Discussion

Phase locking of neural firing to external and internal signals is an ubiquitous and important physiological mechanism in the brain (Köppl, 1997; Surlykke et al., 1988; Joris and Smith, 2008; Taniguchi and Hisashu, 1987; Bastian, 1981; Chacron et al., 2000; Ratnam and Nelson, 2000; Montemurro et al., 2008; Martin and Schröder, 2016; Ewert et al., 2008; Wubbels, 1992; Buzsáki and Draguhn, 2004). It allows neurons to encode the temporal structure of stimuli and decode important stimulus information from minute temporal differences in spike arrivals (Köppl, 1997). Some neurons have intrinsic dynamics that let them resonate at single or multiple frequencies, which allows them to read out periodic phase locked inputs (Llinas, 1988; Hutcheon and Yarom, 2000; Buzsáki and Draguhn, 2004; Neiman and Russel, 2004). Locking in individual neurons is usually studied for single frequencies only. Here, we studied multiple frequency locking in single neurons in the electrosensory system of the weakly electric fish by exploring the entire spectrum of locking frequencies at once (van Hemmen et al., 2011). We identify key elements for multiple frequency locking and provide concise mathematical models that recreate and explain our main findings.

Weakly electric fish extract social and environmental information from modulations of their electric field. In particular the EODs of other wave-type electric fish provide a second, periodic input to electroreceptor afferents resulting in beating AMs. In case of more than two interacting fish, more complicated AMs arise (Stamper et al., 2012). Syntopic species occupy specific, mostly separate EOD frequency ranges (Crampton and Albert, 2006; Kramer et al., 1981), and the EOD frequencies of individual conspecifics differ (Fig. 9, Henninger et al., 2017). In a given habitat, weakly electric fish are therefore exposed to a wide range of EOD frequencies from conspecifics as well as heterospecifics, ranging from about 50 Hz to more than 1000 Hz. Although P-units are tuned to the EOD frequency (Knudsen, 1974; Hopkins, 1976), we observed significant locking of P-unit spikes to all these frequencies (Fig. 3).

**Figure 9.**
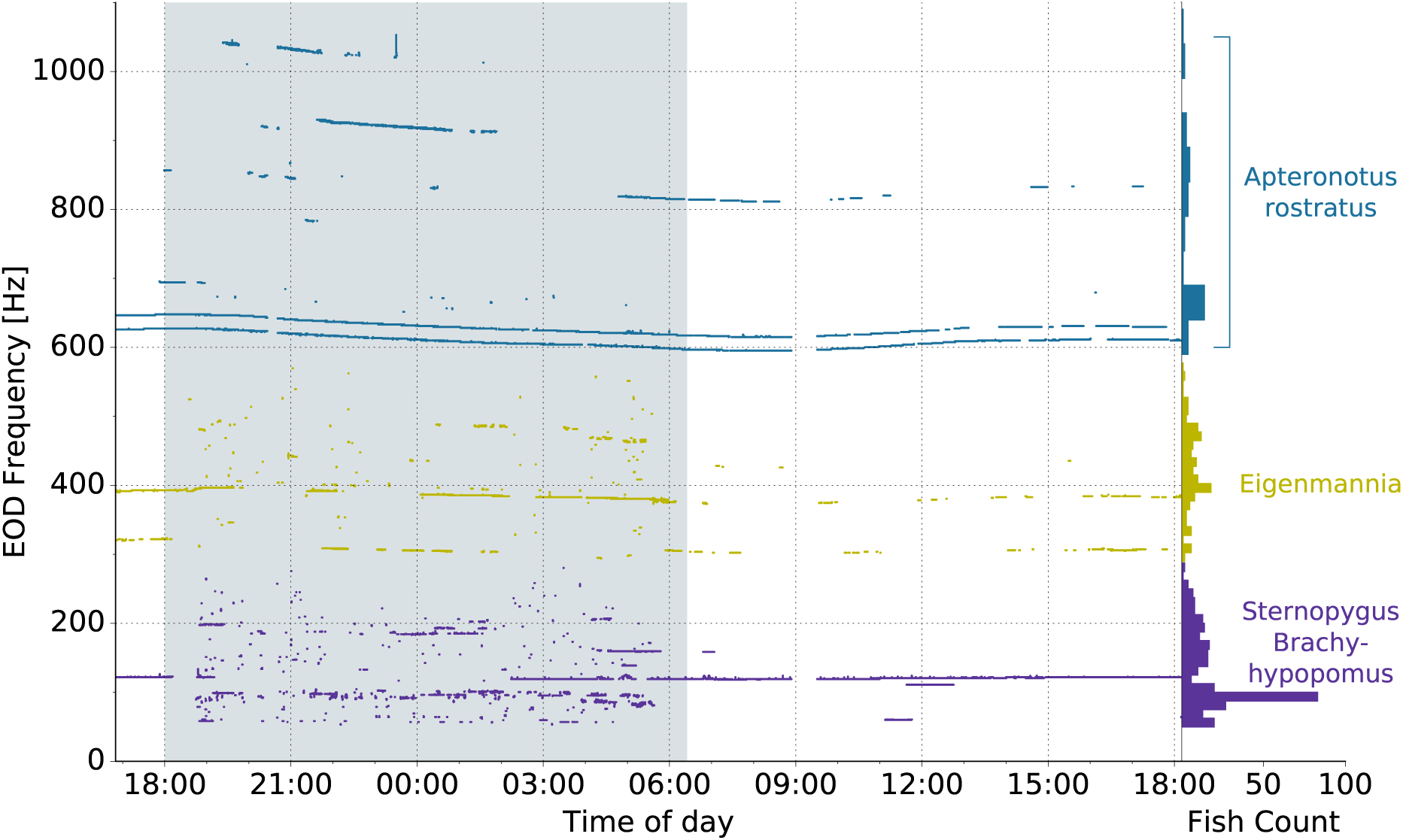
EOD frequencies of weakly electric fish recorded in a single habitat. Recording of electric fish activity in an 54 electrode grid covering 3.6 m^2^ of a neotropical stream in the Province of Darién, Republic of Panamá (see Henninger et al. (2017) for details). Each dot in the main panel marks the fundamental frequency of an observed EOD over a 25 h recording period. Most of the time several fish with distinct EOD frequencies are simultaneously present in the recording area. The shaded area marks the dark period. The histograms at the right show the distribution of observed EOD frequencies of the three wave-type fish *Apteronotus rostratus*, *Eigenmannia humboldtii*, and *Sternopygus dariensis*. The frequencies of the pulse-type fish *Brachyhypopomus occidentalis* below 100 Hz overlap with *Sternopygus*.

The common view is that each of the three different types of specialized electroreceptors afferents in gymnotiform weakly-electric fish encodes a particular signal feature of the electric environment: ampullary electroreceptor afferents for slow external electric fields, T-type electroreceptor afferents (T-units) for phase information, and P-type electroreceptor afferents (P-units) for AMs of the own field (Wessel et al., 1996; Chacron, 2007; Stöckl et al., 2014). In particular, P-units are thought to mainly encode the AM of the fish’s EOD in their spike rate. We demonstrate that they also contain rich phase information and can encode the absolute frequency of external signals over a wide frequency range via phase locking. In fact, their vector strength spectrum shows locking to many frequencies including the EOD, the stimulus, the AM, and harmonic combinations thereof. This multiple locking phenomenon is made possible by the fine temporal structure of P-unit firing which is locking to particular phases of the fish’s own EOD. When their spikes are selectively suppressed in the presence of an external signal, locking to other — in particular higher — frequencies can emerge. This fine temporal structure in their spike pattern would not be needed for encoding the AM in a rate code. In fact, previous work in *E. virescens* has found that spike times can be substantially jittered in time before the decoding performance of the AM decreases (Kreiman et al., 2000). Thus, our findings show that the previous characterization of P-units as rate coders of the AM is too narrow, and that P-units are an example of how rate and temporal codes can coexist and complement each other to provide a rich representation of the sensory environment (Panzeri et al., 2015).

A previous study in *E. virescens* showed that phase information can be decoded from P-units and amplitude information can be decoded from T-units when the preferred stimulus feature of the respective unit was sufficiently weak (Carlson and Kawasaki, 2006). The authors reconstructed random AMs and phase modulations (PMs) of a carrier close to the EOD frequency from the evoked spike trains. While decoding AMs and PMs was possible from both receptor types when the modulations were presented in isolation, the reconstruction quality of the receptor’s non-preferred stimulus modality deteriorated when the two modulations were presented simultaneously. The authors showed that this inherent ambiguity in phase and amplitude information in both receptors can be used to evoke an incorrect jamming avoidance response from the fish (Carlson and Kawasaki, 2006; Heiligenberg, 1991). This demonstrates that phase information in P-units affects the fish’s perception.

Combined with our results, these findings (Carlson and Kawasaki, 2006) demonstrate that P-units are not just simple amplitude encoders, but provide the electric fish with a rich source of information. Further evidence that this source of information is useful and behaviorally relevant is provided by a recent study in *E. virescens* that show that electric communication signals, called *chirps*, can be more reliably decoded from responses of three P-units than from responses of an ampullary electroreceptor afferent, a T-unit, and a P-unit (Stöckl et al., 2014). Chirps have complex effects on all receptor types and one would expect that a more heterogeneous population of receptors provides more information about the chirp type (Stöckl et al., 2014; Benda et al., 2006). The fact that P-units yield such a good decoding performance indicates that they represent an important information resource for the decoding of social signals.

P-units converge on pyramidal cells (Maler, 2009). Spikes of pyramidal cells also directly lock to the stimulus. This indicates that spike timing information from P-units might actually be used in higher brain areas. Since the ELL is the bottleneck for all electrosensory information, stimulus features that are used higher up in the processing hierarchy must first be present in the firing of ELL cells. Pyramidal cell can receive input from up to a thousand P-units (Maler, 2009). Therefore, it may be surprising that locking to the stimulus is still present after integrating so many inputs from spatially extensive receptive fields. If spikes of single P-units would arrive at different phases of the EOD, the locking effect would average out during synaptic integration at the pyramidal cell. However, previous studies found that axons of P-units in *E. virescens* located in the tail are thicker and thus faster compared to axons of cells in the head region (Heiligenberg and Dye, 1982). The authors speculate that the reason for this morphological adaptation is to minimize the temporal discrepancy of spike arrival times caused by global events such as other fish. Our results indicate that a similar principle might be at work in *A. leptorhynchus* and that spike timing in P-units does matter. Given that these fish are usually smaller than 30 cm (de Santana and Vari, 2013), the jitter induced by conduction in myelinated axons along the fish would hardly matter for the accuracy of a rate code of AMs (Kreiman et al., 2000).

Where and how this locking information is decoded beyond the ELL is still unclear. Since neurons in the *torus semicircularis* are sensitive to the differential phase of two signals (Rose and Heiligenberg, 1985, 1986) this brain area is a likely candidate. Because the interactions between fundamental frequencies and harmonics create a complex locking pattern, separating the different causes of the pattern is not trivial. In general, however, locking to particular frequencies could be read out by resonance mechanisms like the one found in our LIF model or other systems (Llinas, 1988; Hutcheon and Yarom, 2000; Buzsáki and Draguhn, 2004).

Here we showed that multiple frequency locking in p-type electroreceptor afferents arises from two simple ingredients, fine temporal locking to an external signal and selective suppression of single cycles as a consequence of the beating AM induced by the interaction of two frequencies. In particular in auditory systems natural stimuli consisting of more than a single sound frequency are also conceivable. Locking of auditory nerve fibers to low-frequency sounds (Köppl, 1997) as well to amplitude modulations (Joris and Yin, 1992) closely resembles response properties of p-type electroreceptor afferents. We therefore expect similar locking spectra and mechanisms in auditory nerve fibers when stimulated with a superposition of two pure tones. Since locking is further improved in cochlear nucleus (Joris and Smith, 2008), the information carried by locking to multiple frequencies might play an important role in the auditory system.

## Methods and Materials

### Electrophysiology

In vivo intracellular recordings of electroreceptor afferents or pyramidal neurons in the Electrosensory lateral line lobe (ELL) were performed on *n* = 9 fish of the brown ghost knifefish *A. leptorhynchus*. All experimental procedures complied with German and European regulations and were approved by the local district animal care committee (file no. ZP 1/13).

#### Surgery

Fish were initially anaesthetized with 150 mg/L MS-222 (buffered to pH of 7.0 using NaHCO_3_, PharmaQ, Fordingbridge, UK) until the deepest level of anaethesia was reached, i.e. gill movements ceased. Fish were then respirated with a constant flow of water through a mouth tube. Respiration water contained 120 mg/L MS-222 (buffered to pH 7.0) during the surgery to sustain anaesthesia. To record from electroreceptor afferents, the lateral line nerve was exposed by cutting a small opening in the skin dorsal to the operculum where the nerve comes very close to the surface before entering the brain. For recordings of pyramidal cells in the hindbrain, the respective parts of the skull were exposed and a hole was drilled into the skull bone. All surgical wounds were locally anaesthetized with liquid Lidocainehydrochloride 2% (bela-pharm GmbH, Vechta, Germany), which was renewed every two hours.

After surgery, fish were immobilized by injecting 0.05ml (1 mg/ml) tubocurarine (Sigma - Aldrich, Steinheim, Germany) into the trunk muscles and respiration was continued without anaesthesia. Fish were then transferred into the recording tank of the setup. The animals were lowered into the water such that the surgical wounds were above the water surface. Respiratory and tank water were taken from the housing tanks to maintain conditions, water temperature was kept at 26° C.

#### Recording and stimulation

Intracellular recordings were done with sharp borosilicate microelectrodes (GB150F-8P, Science Products, Hofheim, Germany), pulled to a resistance between 60 and 100MΩ and filled with a 1 M KCl solution. Electrodes were positioned by microdrives (Luigs-Neumann; Ratingen, Germany). As a reference, chlorided silver wire was used which was placed in the tissue surrounding the exposed nerve or the hole in the skull bone. The potential between the micropipette and the reference electrode was amplified (SEC-05X, npi electronic GmbH; Tamm, Germany) and lowpass filtered at 10 kHz. Signals were recorded by a data acquisition board (PCI-6229, National Instruments, Austin TX, USA) at a sampling rate of 20 kHz. Spikes were detected and identified online according to the algorithm proposed by Todd and Andrews (1999).

The EOD of the fish was measured between the head and tail via two carbon rod electrodes (11 cm long, 8mm diameter). The potential at the skin of the fish was recorded by a pair of silver wires, spaced 1cm apart, which were placed orthogonal to the longitudinal axis of the fish at one thirds body length. These EOD voltages were amplified and bandpass filtered with cutoff frequencies of 3 Hz and 1.5 kHz for low and high frequency cutoff, respectively (DPA-2FXM, npi-electronics, Tamm, Germany).

The EODs of foreign fish were mimicked with pure sinewaves of frequencies in the range −500 to +500 Hz relative to the recorded fish’s EOD. Stimulus amplitudes were adjusted to evoke AMs of certain contrasts. That is, the AM depth relative to the fish’s EOD amplitude was adjusted to have contrasts of 2.5, 5, 10 or 20% at the measurement electrode at one third of the body length (above). Stimuli were attenuated (ATN-01M, npi-electronics, Tamm, Germany), isolated from ground (ISO-02V, npi-electronics, Tamm, Germany), and delivered by two carbon rod electrodes (30cm length, 8mm diameter) placed on either side of the fish parallel to its longitudinal axis. Spike and EOD detection, stimulus generation and attenuation, as well as pre-analysis of the data were performed online during the experiment using the efield and efish plugins of the RELACS software (www.relacs.net).

## Analysis

The analysis was performed using the Python libraries scipy (Oliphant, 2007), numpy (van der Walt et al., 2011), matplotlib (Hunter, 2007), pandas (McKinney, 2010), statsmodels, nix (Stoewer et al., 2014), seaborn (Waskom et al., 2016), and datajoint (Yatsenko et al., 2015). All analysis code is freely available from https://github.com/fabiansinz/efish_locking.git or can be requested from the corresponding author. A Docker container with the pre-installed code can be pulled from https://hub.docker.com/r/fabee/locking/. Data can be requested from the corresponding authors.

### Vector strength spectra and Fourier analysis

Let a spike train be given by its spike times *t_i_* or delta pulses *δ* (*t* − *t_i_*) at the spike times respectively. Then we get the following relation between the amplitude spectrum of the spike train and the vector strength *ν*(*f*) of the spike train relative to the period 1*/f* of a signal with frequency *f* (Ashida et al., 2010):

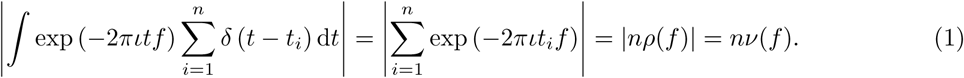

Here, *ρ*(*f*) denotes the average resultant vector of the events at phases *α_i_* = 2*πt_i_f* and *ν*(*f*) = |*ρ*(*f*)| denotes the corresponding vector strength.

### Statistical tests on locking

There are two different ways how several spike trains can enter the estimation of the vector strength spectrum: The vector strength could be computed for each trial and the spectra would be averaged afterwards (second order spectra), or the spikes of all spectra could be aggregated beforehand and the spectrum would be computed afterwards (first order spectra). These two possibilities lead to two different results concerning the statistical information about the spike train they contain and concerning the null distributions for tests against non-locking.

Aggregating the spikes before computing the vector strength for a particular locking frequency is equivalent to taking the Fourier transform of the PSTH in the limit of infinitely many trials. When the spikes do not lock to this particular frequency, the vector strength will converge to zero as the number of trials grows. Statistical significance on the aggregated spike trains can be assessed by Rayleigh’s test (Ashida et al., 2010). However, aggregating spike times before computing the vector strength will remove the per trial dependencies between the spike times, because shuffling spikes across trials results in the same set of aggregated spike times. Thus, the resulting first order vector strength spectrum usually exhibits less peaks than averaging vector strength spectra across trials.

Computing the vector strength per trial and then averaging across trials preserves the per trial interactions between the stimulus and the spikes. However, even when the spike times do not lock to this particular frequency, the average vector strength will not converge to zero with an increasing number of trials as in the former case, but to some value determined by the number of spikes per trial. We discuss this case and associated statistical tests in more detail.

Consider a Poisson null distribution for the spikes used to compute *ν*(*f*). In that case the spike phases have a uniform distribution on the complex circle. The vector strength *ν*(*f*) = |*ρ*(*f*)| is the absolute value of the average vector *ρ*(*f*) of those points. For large enough number of spikes the average of *n* independent realizations of spike times converges to an isotropic Gaussian distribution 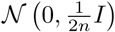 because of multivariate central limit theorem and the spherical symmetry of the uniform distribution on the circle. *I* denotes the identity matrix in the complex plane. Consequently, the radial density of this Gaussian is a *χ*-distribution.

In the aggregated case one can construct Rayleigh’s test from the *χ*-distribution. Since the Gaussian has zero mean and the variance depends inversely on *n*, the sample vector strength will converge to zero with an increasing number of spikes. If, however, the vector strength is computed first and then averaged over trials, the variance of the Gaussian null distribution and therefore the mean of the *χ*-distribution, stays the same per trial. This means that averaging the spectra will not lead to a vector strength of 0 in the limit of infinite trials but converge to a positive number that is determined by the number of spikes per trial.

In order to compute a confidence interval for this case, we start with a single trial of *n* Poisson distributed spikes. Using the central limit theorem as above, the sampling distribution of the vector strength for this spike train is approximately a *χ*-distribution *p*(*r*|*n*) = 2*nr* · exp (−*nr*^2^). This approximation requires the number of spikes to be sufficiently large to make *ρ*(*f*) approximately Gaussian. If the number spikes is Poisson distributed with some rate λ, the sampling distribution of the vector strength *ν*(*f*) becomes a mixture distribution

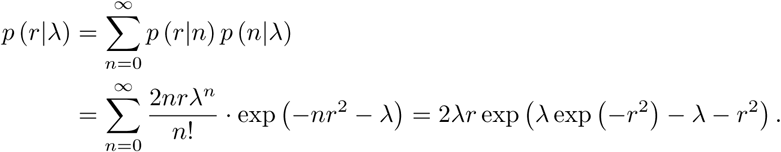

Because *p* (*r*|*n*) is not a density for *n* = 0 we can refine our distribution by exluding this case and rescaling the Poisson distribution. Then then final sampling distribution is given by

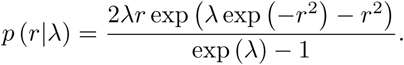

This is the density of the random variable *ν*(*f*) if the number of spikes is drawn from a Poisson distribution. From that expression, we numerically compute the mean and the variance using the empirical estimate 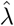 of the firing rate. For *N* trials, we scale the resulting standard deviation by 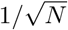 and compute a one-sided confidence interval using a Gaussian distribution with the previously determined mean.

### Phenomenological Model

The phenomenological model explains locking to the external stimulus — and weaker locking to its harmonics — by locking of the P-unit to the EOD and Gaussian jitter on the spike times. If EOD*f* denotes the EOD frequency, we assume that each P-unit fires at a particular phase of the EOD period of length *T* = 1*/*EOD*f* and that the spike times are jittered by Gaussian random noise with variance 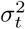 (Fig. 4 C, left). This means that the PSTH is the convolution III*_T_* _\**g*_ of a Dirac comb 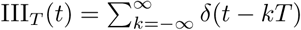 with period *T* with a Gaussian 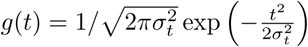. According to the convolution theorem, the vector strength spectrum is therefore proportional to a Dirac comb with period EOD*f* multiplied with a Gaussian with variance 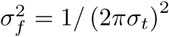 (Fig. 4 C, right).

A simple way to capture the modulation of the firing rate in the presence of a stimulus is to multiply the Dirac comb with *a*cos(2*π*Δ*f*) + *b* before convolving it with a Gaussian. This modulates the unjittered PSTH with the beat frequency (Fig. 4 C-D). In the Fourier domain, this multiplication becomes a convolution with two Dirac peaks at −Δ*f* and Δ*f*. This convolution causes each spike of the Dirac comb to be flanked by two additional peaks (see fig. 4 D, right), as observed in vector strength spectra of our data.

### Model

We modeled spike generation in P-units by a leaky integrate and fire (LIF) model

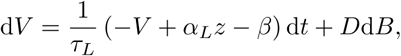

where *V* is the modeled membrane voltage, *τ_L_* denotes the membrane time constant, *α_L_* and *β* model an affine transformation of the input *z*, and d*B* is the differential for Gaussian noise with intensity *D*.

The membrane potential was driven by the output *z* of a rectified, low-pass filtered, damped harmonic oscillator, which was driven by the external electric field *s*:

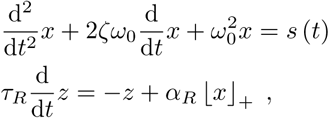

where *ζ* is the damping ratio of the harmonic oscillator and

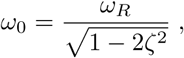

is its undamped angular frequency with *ω_R_* = 2*πf_R_*. The resonant frequency *f_R_* was set to the measured EOD frequency of each fish. *τ_R_* is the time constant of the low-pass filter and its input gain *α_R_* was set to

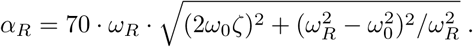

⌊·⌋_+_ denotes rectification. We used *ζ* = 0.2, *τ_R_* = 0.002, *α_L_* = 1, *β* = 9, *D* = 30, and *τ_L_* = 0.001. The spiking threshold was set to 14 and the reset potential was set to 0. The harmonic oscillator was solved using the differential equation solver from SciPy (Oliphant, 2007). The LIF was solved using Euler’s method.

## Acknowledgements

This study is part of the research program of the Bernstein Center for Computational Neuroscience, Tbingen, funded by the German Federal Ministry of Education and Research (BMBF; FKZ: 01GQ1002). Fabian Sinz was supported by a grant of the German Research Foundation (DFG) awarded to Jan Benda (CRC 870). Carolin Sachgau was supported by the Ontario/Baden-Wrttemberg Summer Research Program. The authors would like to thank Philipp Berens for comments on the manuscript.

